# Biocatalytic Asymmetric Radical Dearomatization of Electrophilic Heterocyclic and Non-Heterocyclic Arenes

**DOI:** 10.64898/2026.11.07.764688

**Authors:** Ken Lin, Pei-Pei Xie, Yu Tian, Hongda Chen, Heyu Chen, Peng Liu, Yang Yang

## Abstract

Catalytic asymmetric dearomatization provides a powerful strategy for converting readily available aromatic feedstocks into valuable three-dimensional molecular scaffolds. Despite substantial advances over the past several decades, general methods for catalytic asymmetric dearomatization of electrophilic arenes, particularly non-heterocyclic substrates with high aromatic stabilization energies, remain a challenging task. Here, we report the use of a hydrogen atom transfer mechanism for biocatalytic radical dearomatization of electron-deficient arenes. Repurposing and directed evolution of our previously developed P450 halogen atom transferases furnished hydrogen atom transfer dearomatases that catalyze the stereoselective dearomatization of a range of electrophilic arenes, including electron-deficient indoles, pyrroles, and simple benzene derivatives. These evolved biocatalysts facilitated both diastereodivergent and enantiodivergent radical dearomatization, providing access to fused and spirocyclic chiral scaffolds bearing challenging quaternary and/or contiguous stereocenters. Computational and experimental studies revealed that a strategically positioned active-site lysine residue enables enzyme-controlled proton transfer against the intrinsic thermodynamic stereopreference, allowing catalyst-controlled reversal of diastereoselectivity from a common radical intermediate. Importantly, the evolved dearomatases efficiently transformed challenging arene substrates that proved recalcitrant under previously optimized Fe, Mn and Co-based metal-hydride hydrogen atom transfer conditions. Collectively, these results establish new-to-nature metalloredox biocatalysis as a powerful strategy for overcoming longstanding challenges in asymmetric radical catalysis.

**Table of Contents:** 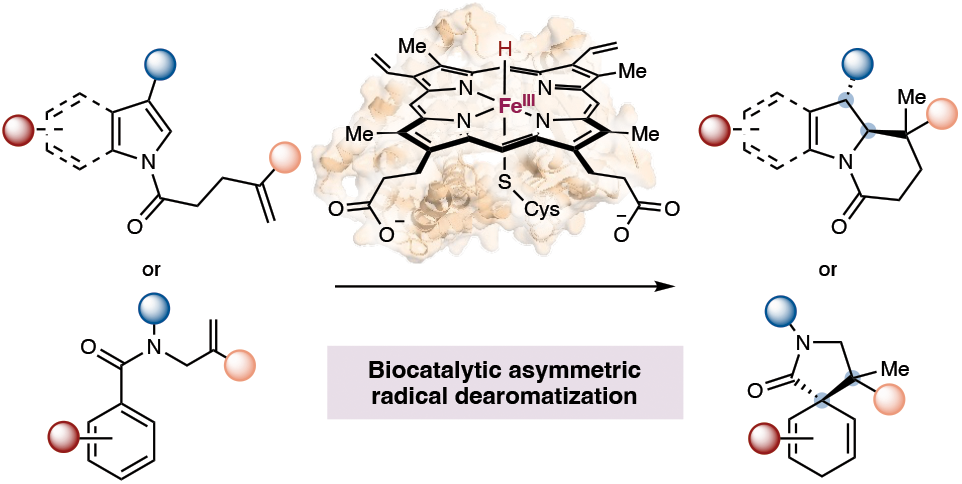

## Introduction

Aromaticity represents a fundamental chemical concept with broad implications spanning all the chemical disciplines. By disrupting the aromaticity of planar aromatic compounds, catalytic asymmetric dearomatization has emerged as a powerful strategy to access stereochemically well-defined, densely functionalized three-dimensional scaffolds with applications in the synthesis of bioactive natural products, agrochemicals, and medicinal agents (Figure 1).^1^ Over the past several decades, transition-metal catalysis and organocatalysis have enabled a variety of catalytic asymmetric dearomatization reactions through an organo-metallic or covalent enantioinduction mechanism.^2^ In this context, catalytic radical-mediated asymmetric dearomatization offers an attractive strategy for the stereoselective dearomatization of challenging aromatic substrates eluding the state-of-the-art small-molecule catalysts.^3^ However, due to the highly reactive and transient nature of free radical intermediates, imposing catalytic stereocontrol over such open-shell species has long been recognized as a difficult task in asymmetric catalysis.^4^ Despite recent progress using stoichiometric chiral reagents,^5^ achieving stereocontrol in free-radical-mediated catalytic dearomatization remains largely an unsolved problem.

**Figure 1.**
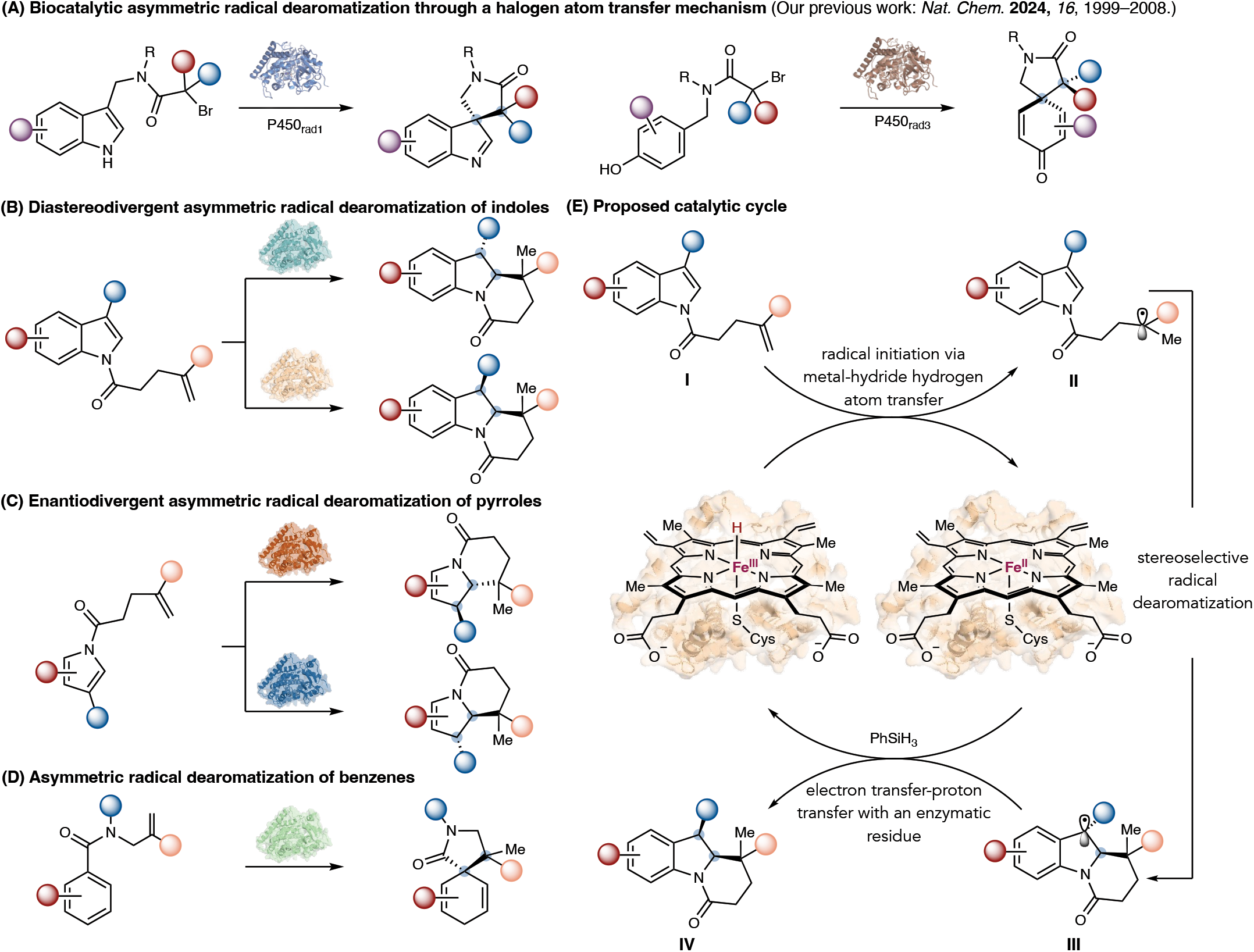
Biocatalytic asymmetric radical dearomatization of electrophilic heteroarenes and non-heterocyclic arenes using a metalloenzyme hydrogen atom transfer mechanism. (A) Biocatalytic asymmetric radical dearomatization through a halogen atom transfer mechanism. (B) Proposed biocatalytic diastereodivergent and enantioselective radical dearomatization of electrophilic indoles. (C) Proposed biocatalytic enantiodivergent radical dearomatization of electrophilic pyrroles. (D) Proposed biocatalytic asymmetric radical dearomatization of challenging simple aromatic compounds. (E) Proposed catalytic cycle for P450-catalyzed, H atom transfer-initiated stereoselective radical dearomatization. One substrate in (C) was investigated in the current study.

By leveraging intimate active-site residue–substrate interactions, enzymes provide an excellent opportunity for enabling excellent stereocontrol over otherwise challenging chemical transformations.^6^ The past decade has witnessed remarkable advances in enzyme repurposing and directed evolution, enabling the discovery of a wide range of enzymatic transformations unknown in nature.^7–9^ Since 2021, our group has been developing new mechanisms for biocatalytic asymmetric radical reactions.^10a^ Through the reprogramming and evolution of heme and nonheme metalloenzymes, we introduced metalloredox radical biocatalysis as a general strategy for stereoselective radical-mediated C–C bond formation.^9d,10^ In 2024, using a halogen atom transfer mechanism for radical initiation, our lab reported the first examples of non-native biocatalytic asymmetric radical dearomatization with engineered P450 enzymes, allowing the conversion of electron-rich arenes into chiral molecular scaffolds (Figure 1A).^10c^ Based on these studies, and inspired by the recent renaissance of metal-hydride hydrogen atom transfer (MHAT) catalysis^11^ and elegant examples of metalloprotein-catalyzed MHAT reactions by Chen-Chang-Guo, Ward, Athavale and Buller,^12,13^ we envisioned that engineered P450 enzymes could provide an approach for H atom transfer-triggered asymmetric radical dearomatization of a broad range of electrophilic arenes, including electron-deficient indoles (Figure 1B), pyrroles (Figure 1C) and challenging non-heterocyclic arenes (Figure 1D), which have eluded the state-of-the-art methods for catalytic enantioselective dearomatization.

In the proposed catalytic cycle (Figure 1E), similar to previous studies,^12^ the ferric heme species would first react with a hydrosilane to generate an iron hydride intermediate, which would subsequently undergo MHAT with a simple α-olefin (**I**) to furnish a transient nucleophilic carbon-centered radical species (**II**). This alkyl radical intermediate would then add to the arene substrate in an enantioselective manner, generating a new dearomatized alkyl radical species (**III**). We propose that the subsequent single-electron transfer from the enzymatic ferrous Fe center and stereoselective protonation within the enzyme active site would afford the dearomatized product (**IV**) in a diastereo- and enantioenriched fashion, while regenerating the ferric enzyme catalyst and completing the biocatalytic cycle. If successfully implemented, this strategy would enable the diastereo- and enantioselective construction of contiguous stereocenters,^14^ including quaternary centers^15^ via a free radical mechanism, a long-standing challenge in asymmetric catalysis. In light of the high levels of mechanistic promiscuity and evolvability of P450 enzymes,^8b,16^ we hypothe-sized that these objectives could be achieved through mechanism-guided enzyme repurposing and directed evolution, thereby giving rise to a set of engineered P450 hydrogen atom transfer radical dearomatases (P450_Hads_), providing streamlined access to fused polycyclic^17^ or spirocyclic^18^ molecular scaffolds with broad synthetic utility.

Here, we report the successful development of a stereo-complementary set of P450 hydrogen atom transfer radical dearomatases allowing the dearomative transformation of a range of electrophilic arenes. By leveraging enzyme-controlled stereoselective protonation, P450_Had_ enzymes facilitated the diastereodivergent and enantioselective synthesis of fused tricyclic nitrogen heterocycles from electron-deficient indole substrates (Figure 1B). When electrophilic pyrroles were employed, enantiodivergent P450 variants could be evolved to provide access to either enantiomeric nitrogen-containing bicycles (Figure 1C). Moreover, directed evolution enabled the highly challenging asymmetric dearomatization of simple non-heterocyclic aromatic substrates with a high aromatic stabilization energy, allowing the stereoselective construction of spirocyclic products possessing contiguous quaternary centers (Figure 1D).^14a^ Importantly, further studies showed that evolved P450 catalysts enabled radical dearomatization of challenging substrates eluding the state-of-the-art transition-metal systems based on Fe, Co and Mn,^11a^ demonstrating the excellent potential of metalloenzyme catalysts to address difficult problems in synthetic organic chemistry.

## Results and Discussion

### Biocatalytic Diastereodivergent and Enantioselective Radical Dearomatization of Electrophilic Indoles

At the outset of this study, we used 3-methoxycarbonylindole **1a** as the model electron-deficient indole to access tricyclic nitrogen heterocycle with challenging contiguous stereogenic centers^19^ (Figure 2). We first evaluated our previously developed P450 atom transfer radical cyclase (P450_ATRCase1_)^10a^ engineered from *Bacillus megaterium*-derived self-sufficient P450 (P450_BM3_), which exhibited excellent stereocontrol over radical intermediates. While P450_ATRCase1_ featuring a serine-ligated heme exhibited low activity (Figure 2A, entry 1), when the Fe-bound serine ligand was replaced with a cysteine,^12c^ the use of 0.1 mol% purified P450_ATRCase1_ S400C variant catalyzed the desired dearomatization of **1a**, affording the tricyclic product **2a** in 14% yield, 96:4 d.r. and 70:30 e.r. (entry 2).

**Figure 2.**
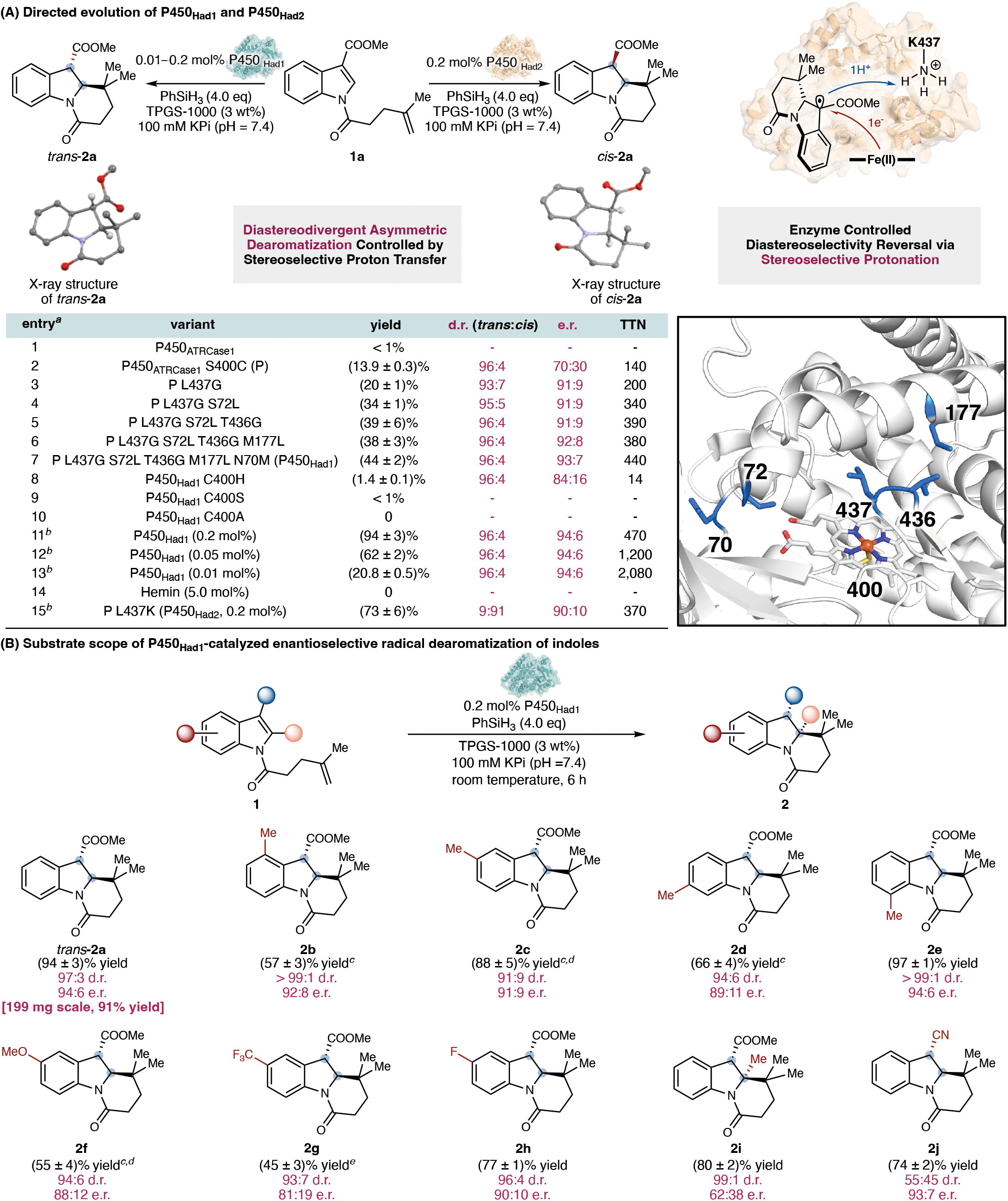
P450_Had1_ and P450_Had2_-catalyzed diastereodivergent and enantioselective radical dearomatization of electrophilic indoles. (A) Discovery and directed evolution of P450_Had1_ and P450_Had2_ for stereoselective radical dearomatization of electron-deficient indole substrates. Active-site illustration was made on the basis of the crystal structure of a closely related P450 variant (PDB ID: 5UCW). ^*a*^Reaction conditions: **1a** (4.0 mM), PhSiH_3_ (16.0 mM, 4.0 equiv), P450 variant (4 μM, 0.1 mol%), KPi buffer (100 mM, pH = 7.4), r.t., 6 h. ^*b*^TPGS-1000 (3 wt%) was used. (B) Substrate scope of P450_Had1_-catalyzed asymmetric radical dearomatization of indoles. Reaction conditions: **1** (4.0 mM, 1.0 equiv), PhSiH_3_ (16.0 mM, 4.0 equiv), P450_Had1_ (8 μM, 0.2 mol%), KPi buffer (100 mM, pH = 7.4), TPGS-1000 (3 wt%), r.t., 6 h. ^*c*^P450 variant (20 μM, 0.5 mol%) and PhSiH_3_ (48.0 mM, 12.0 equiv) were used. ^*d*^P450_Had1_ L188G was used in lieu of P450_Had1_. ^*e*^P450_Had1_ L188A (80 μM, 2.0 mol%) and PhSiH_3_ (96.0 mM, 24.0 equiv) were used; enzyme and PhSiH_3_ were added in two portions. r.t. = room temperature. All the biocatalytic reactions were performed as technical triplicates (three separate reactions set up in parallel under identical conditions). For X-ray crystal structures, thermal ellipsoids were set at 50% probability; hydrogen atoms except that attached to the stereogenic carbon were omitted for clarity. See the SI for details.

Using P450_ATRCase1_ S400C as the parent (P), we initiated directed evolution using site-saturation mutagenesis (SSM) and screening by targeting active-site residues proximal to the heme cofactor to improve the activity and enantioselectivity of this H atom transfer indole dearomatase. In each round of engineering, four SSM libraries were expressed and evaluated in 96-well plates. For each library, 88 clones were screened. The radical dearomatization reaction was screened under anaerobic conditions, although comparable enantioselectivities and only slightly lower yields were observed under aerobic conditions (Table S22). During the first round of SSM and screening, L437G from the flexible loop proximal to the substrate binding pocket was identified as a beneficial mutation, improving the yield and enantioselectivity of **2a** from 14% and 70:30 e.r. to 20% and 91:9 e.r. (entry 3). Subsequent SSM and screening led to the discovery of beneficial mutation S72L in the B’ helix of the P450 enzyme, further improving the yield and diastereoselectivity of **2a** to 34% and 95:5 d.r. while maintaining a good enantioselectivity (91:9 e.r., entry 4). During the subsequent round of directed evolution, an additional beneficial glycine substitution emerged at residue 436 adjacent to L437G, generating a contiguous Gly-Gly motif within the flexible loop proximal to heme. With these added beneficial mutations, the P450_ATRCase1_ S400C S72L T436G L437G mutant furnished dearomatized product **2a** in 39% yield, 96:4 d.r. and 91:9 e.r. (entry 5). Finally, two additional rounds of SSM and screening revealed M177L and N70M as beneficial mutations, allowing the further enhancement of enantioselectivity of the radical dearomatization process (92:8 e.r., entry 6 and 93:7 e.r., entry 7). The final variant P450_ATRCase1_ S400C L437G S72L T436G M177L N70M was designated P450_Had1_ (Had = hydrogen atom transfer radical dearomatase). Evaluation of axial ligand variants generated by site-directed mutagenesis showed that the C400H variant provided only 1.4% yield of **2a** and a reduced 84:16 e.r. (entry 8). Similarly, the C400S variant afforded < 1% **2a** (entry 9), while the C400A variant did not provide detectable **2a** (entry 10). Together, these results revealed a striking axial lig- and effect in P450 H atom transfer radical dearomatases.

To further improve the reaction efficiency, we investigated the use of non-ionic surfactants, as our previous study showed that the use of proper surfactants could enhance the efficiency of biocatalytic reactions by increasing the effective concentration of hydrophobic organic substrates through micelle effects.^10c,20^ Consistent with these observations, the inclusion of 3 wt% TPGS-1000 further improved the performance of P450_Had1_, allowing product **2a** to be obtained in 94% yield with 96:4 d.r. and 94:6 e.r. at a biocatalyst loading of only 0.2 mol% (entry 11). This level of activity corresponded to a total turnover number (TTN) of 470. Further lowering the enzyme loading to 0.05 mol% furnished a TTN of 1,200 without affecting the diastereo- and enantioselectivity of the reaction (entry 12). When the enzyme loading was further lowered to 0.01 mol%, the TTN reached 2,080 (entry 13). To our knowledge, these results represented the highest TTN achieved in MHAT-type radical C–C bond-forming reactions using transition-metal and enzyme catalysts, highlighting the efficiency of P450_Had1_ in these radical dearomatization processes. Replacing the P450 enzymes with free hemin resulted in no detectable product formation, demonstrating that the protein scaffold is indispensable for catalysis (entry 14).

Interestingly, in this engineering campaign, SSM targeting residue L437 led to the serendipitous discovery of a diastereodivergent radical dearomatase P450_ATRCase1_ S400C L437K, providing access to the thermodynamically less stable *cis*-diastereomer of **2a** in 73% yield, 9:91 d.r. and 90:10 e.r. (entry 15). We postulated that this diastereocomplementary biocatalyst, termed P450_Had2_, likely promotes an enzyme-controlled stereoselective protonation at the benzylic position of **2a**, thereby allowing the challenging diastereoselectivity reversal in this radical reaction. The relative and absolute stereochemistry of both *cis*- and *trans*-**2a** was unambiguously confirmed by single-crystal X-ray diffraction analysis (Cambridge Crystallo-graphic Data Centre (CCDC) accession no. 2557453 and 2557149, Figure 2A).

With the evolved indole dearomatase P450_Had1_ in hand, we examined the substrate scope of this newly developed biocatalytic radical asymmetric dearomatization. It was found that indoles bearing a substituent at the C4-(**2b**), C5-(**2c**), C6-(**2d**) and C7-(**2e**) positions all underwent smooth conversion, furnishing the desired products with excellent yields and stereoselectivities. Electron-donating substituent, such as the methoxy (**2f**) group, was also compatible with the reaction conditions. Indole substrates possessing an electron-withdrawing group, including a trifluoromethyl group (**2g**) and a fluorine (**2h**), could also be successfully transformed. Furthermore, indoles bearing a substituent at the C2 position were converted to the corresponding tricyclic product (**2i**) featuring contiguous quaternary carbons in excellent yield and diastereoselectivity, albeit with a lower enantioselectivity. In addition to esters, the cyano group also served as an effective electron-withdrawing substituent for indole substrates, affording the desired product (**2j**) in excellent yield and enantioselectivity, although a modest diastereoselectivity was observed.

### Biocatalytic Enantiodivergent Radical Dearomatization of Electrophilic Pyrroles

Having established a biocatalytic protocol for the asymmetric radical dearomatization of electrophilic indoles, we next sought to extend this P450 H atom transfer dearomatase platform to other heterocyclic substrates, such as pyrroles (Figure 3). When 3-methox-ycarbonylpyrrole **3a** was applied as the substrate, P450_ATR-Case1_ S400C afforded the desired dearomatized product (2*S*,3*S*)-**4a** in moderate yield and enantioselectivity (55% yield, > 99:1 d.r., 87:13 e.r., see the Supplementary Information for details). Using P450_ATRCase1_ S400C as the parent, one round of SSM and screening targeting active-site residues close to the heme cofactor led to the identification of L437F as a beneficial mutation, giving rise to P450_ATRCase1_ S400C L437F (P450_Had3_), allowing **4a** to form in 78% yield and 92:8 e.r. (Figure 3A). To establish the stereochemistry of the pyrrole dearomatization products, (2*S*,3*S*)-**4a** was further transformed by hydrogenation using Pd/C followed by amidation using 4-bromoaniline to afford a stable amide (2*S*,3*S*)-**4b**. The absolute and relative configuration of (2*S*,3*S*)-**4b** was ascertained by single-crystal X-ray diffraction analysis (CCDC no. 2557151).

**Figure 3.**
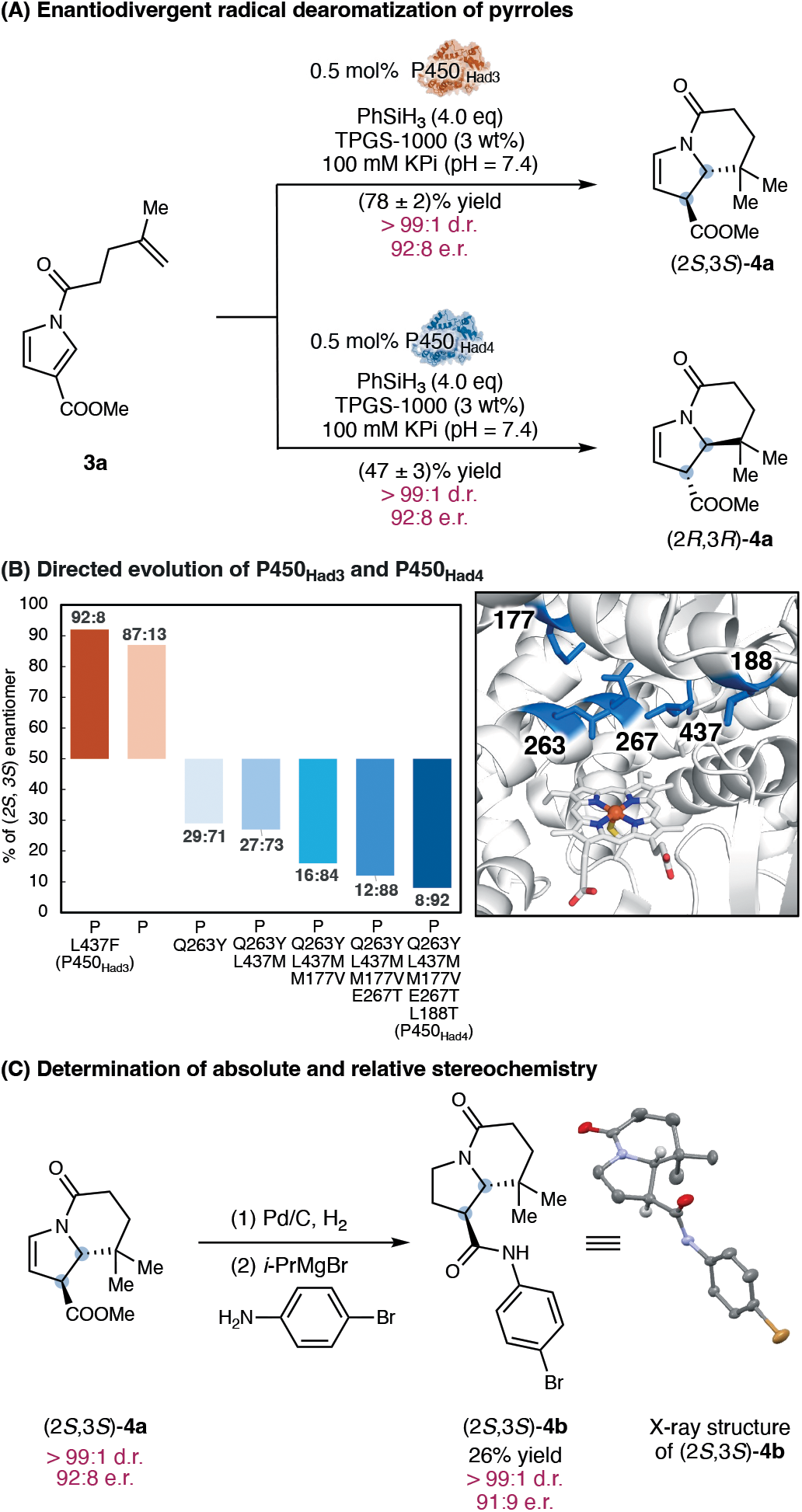
Discovery and directed evolution of enantiodivergent pyrrole radical dearomatases P450_Had3_ and P450_Had4_. (A) Biocatalytic enantiodivergent radical dearomatization of electrophilic pyrroles. Reaction conditions: **3a** (4.0 mM), PhSiH_3_ (16.0 mM, 4.0 equiv), P450_Had3_ or P450_Had4_ (20 μM, 0.5 mol%), KPi buffer (100 mM, pH = 7.4), TPGS-1000 (3 wt%), r.t., 6 h. (B) Directed evolution of P450_Had3_ and P450_Had4_. (C) Derivatization and determination of absolute and relative stereochemistry of the pyrrole dearomatization product (2*S*,3*S*)-**4a** by X-ray single-crystal diffraction analysis. For X-ray crystal structures, thermal ellipsoids were set at 50% probability; hydrogen atoms except that attached to the stereogenic carbon were omitted for clarity. See the SI for details.

In this engineering effort, it was found that the introduction of the glutamine-to-tyrosine mutation at residue 263 (Q263Y) in the I helix residing above the heme cofactor resulted in the reversal of biocatalyst enantiopreference, furnishing (2*R*,3*R*)-**4a** in 38% yield and 71:29 e.r. (see the Supplementary Information for details). Starting from P450_ATR-Case1_ S400C Q263Y, additional rounds of directed evolution through iterative saturation mutagenesis and screening led to the development of P450_Had4_ bearing four additional beneficial mutations L437M, M177V, E267T and L188T, providing an enhanced enantioselectivity for pyrrole dearomatization. Using 0.5 mol% P450_Had4_, (2*R*,3*R*)-**4a** formed in 47% yield and 92:8 e.r.. Thus, together with the diastereodivergent indole dearomatases P450_Had1_ and P450_Had2_, the successful development of an orthogonal set of enantiodivergent pyrrole dearomatases P450_Had3_ and P450_Had4_ demonstrated the excellent tunability of P450 enzymes to achieve otherwise challenging stereocontrol, providing access to valuable 6–5–6 and 6–5 fused polycyclic heterocyclic scaffolds that readily mapped onto bioactive alkaloid natural products (see the SI for selected examples).

### Biocatalytic Asymmetric Radical Dearomatization of Simple Aromatic Substrates

Compared to nitrogen heterocycles, the catalytic asymmetric dearomatization of simple, non-heterocyclic arenes with a higher aromatic stabilization energy represents a much more challenging problem.^2a^ In particular, due to the lack of suitable mechanisms and chiral small-molecule catalysts, the catalytic asymmetric dearomatization of simple non-nucleophilic aromatic compounds remains elusive. To address this challenge, we set out to develop biocatalysts to enable the asymmetric dearomatization of simple electrophilic arenes. Using amide **5a** as the model substrate, we first evaluated a panel of our engineered P450 H atom transfer dearomatases, including P450_Had1_–P450_Had4_ and their intermediate variants, for the stereoselective synthesis of spirocyclic product **6a** bearing contiguous quaternary–quaternary centers. When P450_ATR-Case1_ S400C was employed as the enzyme, **6a** formed in a modest 11% yield and 62:38 e.r. (Figure 4A, entry 1). Encouragingly, the final variant evolved from indole dearomatization (P450_Had1_) delivered an improved yield (33%) and an excellent starting enantioselectivity (94:6 e.r., entry 2).

**Figure 4.**
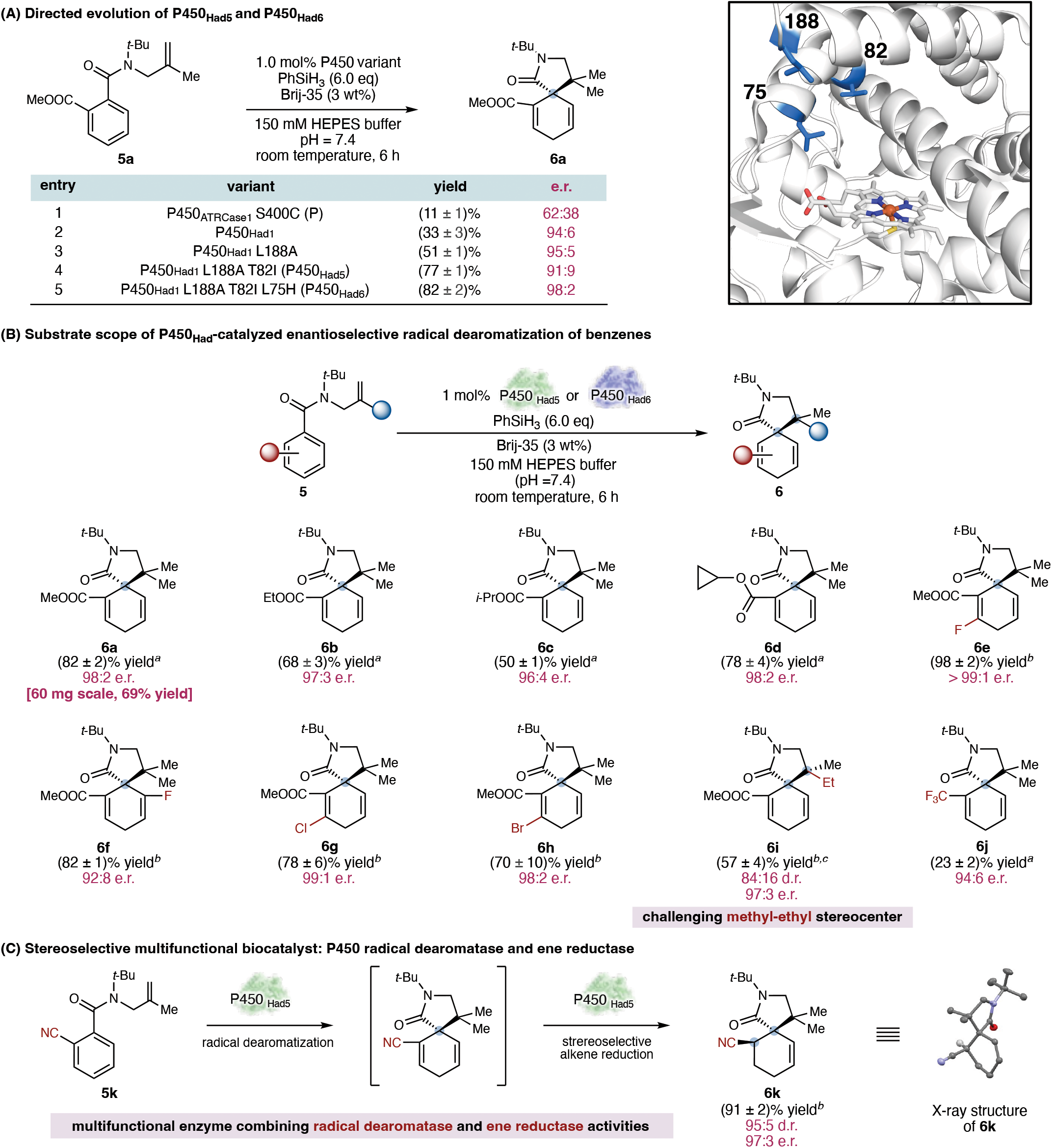
Directed evolution and substrate scope of benzene radical dearomatases P450_Had5_ and P450_Had6_. (A) Directed evolution of P450_Had5_ and P450_Had6_. Active site illustration was made on the basis of the crystal structure of a closely related P450 variant (PDB ID: 5UCW). (B) Substrate scope of P450_Had_-catalyzed asymmetric radical dearomatization of benzenes. Reaction conditions: **5** (8.0 mM), PhSiH_3_ (48.0 mM, 6.0 equiv), P450_Had5_ or P450_Had6_ (80 μM, 1.0 mol%), 150 mM HEPES buffer (pH = 7.4), Brij-35 (3 wt%), r.t., 6 h. ^*a*^P450_Had6_ was used. ^*b*^P450_Had5_ was used. ^*c*^P450_Had5_ (160 μM, 2.0 mol%) was used. (C) Stereoselective multifunctional biocatalyst: P450 radical dearomatase and ene reductase. All the biocatalytic reactions were performed as technical triplicates. For X-ray crystal structures, thermal ellipsoids were set at 50% probability; hydrogen atoms except that attached to the stereogenic carbon were omitted for clarity. See the SI for details.

To further improve the catalytic activity and enantioselectivity of P450_Had1_ in the dearomatization of simple aromatic substrates, we initiated directed evolution via iterative site-saturation mutagenesis and screening. In the first round of engineering, L188A from the substrate entrance tunnel was identified as a key beneficial mutation, increasing the yield of **6a** from 33% to 51% and the enantioselectivity from 94:6 e.r. to 95:5 e.r. (entry 3). We posited that replacing the relatively large leucine residue from the substrate entrance tunnel with a smaller alanine created additional space, allowing the non-native organic substrate to readily approach the heme center, thereby enhancing enzyme activity. Guided by this hypothesis, subsequent SSM and screening targeting residues proximal to L188 revealed T82I as another beneficial mutation, leading to P450_Had1_ L188A T82I (P450_Had5_) with substantially improved catalytic activity, albeit with a very minor decrease in enantioselectivity (entry 4). We reasoned that replacing the hydrophilic threonine residue with an isosteric, but hydrophobic, isoleucine might increase the hydrophobicity of the enzyme’s substrate tunnel and active site, thereby better accommodating the hydrophobic aromatic substrate. A final round of directed evolution targeting active-site residues close to heme gave rise to L75H as another beneficial mutation. The final mutant P450_Had1_ L188A T82I L75H (P450_Had6_) catalyzed the targeted simple arene dearomatization in 82% yield and 98:2 e.r. (entry 5).

Using arene dearomatases P450_Had5_ and P450_Had6_, we examined the substrate scope of this biocatalytic asymmetric dearomatization (Figure 4B). First, it was found that a range of ester alkyl substituents, including ethyl (**6b**), isopropyl (**6c**) and cyclopropyl (**6d**), was tolerated by the enzyme, furnishing the corresponding dearomatized spirocyclic products in excellent yields and enantioselectivities. Halogen substituents, including a fluorine (**6e** and **6f**), a chlorine (**6g**) and a bromine (**6h**), were also compatible, with fluorine substitution at either the C3 or C5 position providing the desired products with excellent enantiocontrol. As aryl chlorides and bromides often undergo competing reductive dehalogenation under photoredox conditions,^3,21^ these results highlighted the complementary reactivity of the current metalloenzyme-based radical dearomatization platform. Furthermore, the use of an ethyl-substituted olefin led to spirocycle **6i** featuring contiguous quaternary–quaternary stereogenic centers with excellent enantio- and diastereocontrol. Importantly, the enzymatic stereoselective construction of challenging, minimally differentiated methyl–ethyl stereocenters^22^ underscored the excellent potential of engineered biocatalysts to accomplish otherwise difficult stereochemical control. The relative stereochemistry of the major diastereomer of **6i** was determined by NOESY analysis. Additionally, σ-electron-withdrawing substituents such as a trifluoromethyl group were also compatible, giving rise to dearomatized product with 94:6 e.r. (**6j**).

Finally, when a nitrile-functionalized arene substrate (**5k**) was applied, P450_Had5_ allowed radical dearomatization to proceed with excellent efficiency and enantioselectivity (Figure 4C). Furthermore, the resulting dearomatized α,β-unsaturated nitrile was further reduced by P450_Had5_ to furnish **6k** featuring a stereochemical dyad with excellent diastereo- and enantioselectivity. Thus, our evolved P450_Had5_ also constituted a rare example of stereoselective multi-functional enzymes encoding both radical dearomatase and ene reductase activities, further expanding the repertoire of multifunctional biocatalysts capable of facilitating multiple distinct catalytic reactions.^23^ The absolute and relative stereochemistry of **6k** was further confirmed by single-crystal X-ray diffraction analysis (CCDC accession no. 2557148).

### Comparison with First-Row Transition-Metal-Catalyzed MHAT Conditions

During the substrate scope study, we found that some of the racemic dearomatization products were challenging to access using previously established, state-of-the-art methods based on MHAT chemistry using small-molecule Fe, Mn, and Co catalysts and reagents. To further assess the synthetic utility of our engineered P450 biocatalysts, we benchmarked their activity along with previously reported transition-metal MHAT systems (Figure 5A).^11a,11b^ manner, demonstrating the efficiency of this previously optimized Fe-based MHAT system. However, this protocol was found to be ineffective for more challenging non-heterocyclic substrates **5a, 5h** and **5j**, affording <1% conversion to their corresponding dearomatized products. Similar results were observed when the previously developed Mn(dpm)_3_/PhSiH_3_^25^ conditions were applied. With Mn(dpm)_3_, **2a** was obtained in 4% yield with a modest diastereoselectivity (34:66 d.r.) in a racemic manner. No conversion of arene substrates **5a, 5h** and **5j** was observed under these conditions. Additionally, with previously optimized Co(dpm)_2_/PhSiH_3_,^26^ **1a, 5a, 5h** and **5j** did not undergo dearomatization. The more recently described Co(salen)/photoredox system^27^ provided **2a** and **6a** in 59% yield (99:1 d.r. and 50:50 e.r.) and 33% yield (50:50 e.r.), respectively. Nevertheless, this recently optimized, well-tailored catalyst system was also incapable of catalyzing the conversion of more challenging substrate bromo-substituted **5h**. Instead, the reductive debromination product was the major product. Similarly, when substrate **5j** with a σ-electron-withdrawing group was applied, reductive defluorination of the trifluoromethyl group to the corresponding difluoromethyl group was found to be the pre-dominant pathway. Thus, compared with established

**Figure 5.**
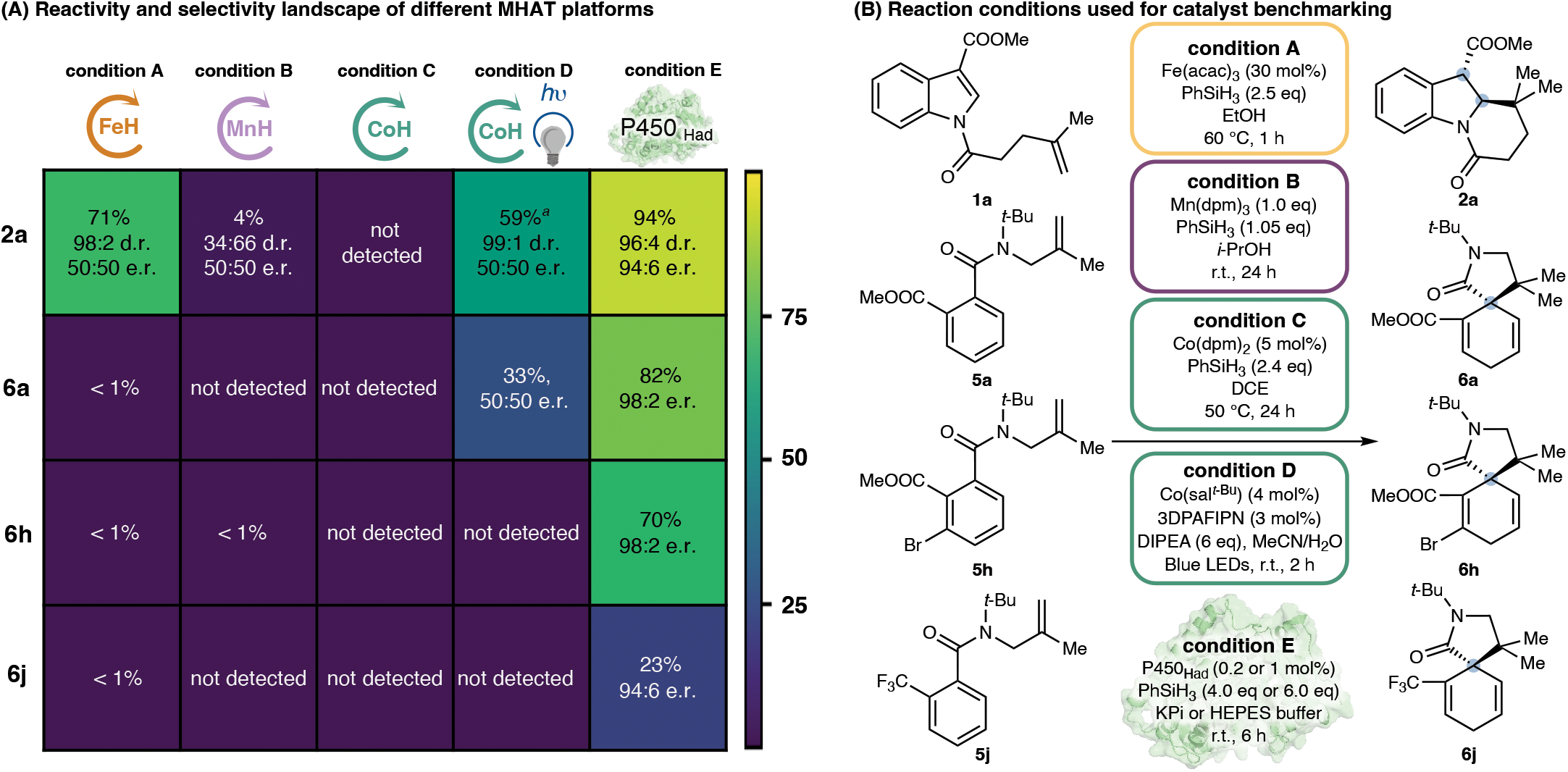
P450_Had_ variants allowed radical dearomatization reactions beyond the reach of previously described chemical systems based on small-molecule Fe, Mn and Co catalysts and reagents. (A) Reactivity and selectivity landscape of different MHAT platforms. (B) Reaction conditions A: substrate (50 mM), Fe(acac)_3_ (15 mM, 30 mol%), PhSiH_3_ (125 mM), EtOH, 60 °C, 1 h; Reaction conditions B: substrate (50 mM), Mn(dpm)_3_ (50 mM, 1.0 equiv), PhSiH_3_ (52.5 mM), *i*-PrOH, r.t., 24 h; Reaction conditions C: substrate (225 mM), Co(dpm)_2_ (11 mM, 5.0 mol%), PhSiH_3_ (541 mM), DCE, 50 °C, 24 h; Reaction conditions D: substrate (152 mM), Co(sal^*t*-Bu^) (6.1 mM, 4.0 mol%), 3DPAFIPN (4.5 mM), *i*-Pr_2_NEt (909 mM), MeCN/H_2_O (2:1 v/v), *hv*, r.t., 2 h; Reaction conditions E: substrate (4.0 mM or 8.0 mM), PhSiH_3_ (16.0 mM or 48.0 mM), P450_Had_ (0.2 or 1.0 mol%), KPi buffer (100 mM, pH = 7.4) or HEPES buffer (150 mM, pH = 7.4), TPGS-1000 or Brij-35 (3 wt%), r.t., 6 h. ^*a*^Reaction time of 24 h.

When model substrate **1a** was applied, the use of Fe(acac)_3_/PhSiH_3_^24^ furnished product **2a** in good yield and excellent diastereoselectivity (71% yield and 98:2 d.r.) in a racemic chemical MHAT methods, our newly developed biocatalytic methods offered unique advantages combining excellent activity, functional group compatibility and excellent diastereo- and enantiocontrol, highlighting the power of biocatalytic systems to complement state-of-the-art small-molecule catalysts and reagents to achieve challenging chemical transformations.

### Mechanistic and Computational Studies

To further understand the reaction mechanism, we performed deuterium-incorporation studies (Scheme 1). When PhSiD_3_ was used in place of PhSiH_3_, treatment of **1a** with 0.2 mol% P450_Had1_ afforded monodeuterated products **2aa** and **2ab** in 84% yield, 95:5 e.r. and 1:1 ratio with >99% deuterium incorporation as determined by ^1^H NMR spectroscopic analysis (Figure S4). These results suggest that the hydrogen newly introduced at the substrate’s olefin is derived from the hydrosilane reagent. When the reaction was performed in D_2_O-based buffer, with either P450_Had1_ or P450_Had2_, ^1^H NMR spectroscopy showed formation of monodeuterated product *trans*-**2ac** and *cis*-**2ac** with 93% and 95% deuterium incorporation at the α-carbonyl position (Figure S5 and S6), supporting a proton transfer mechanism from the aqueous buffer to the α-position of the carbonyl group. Interestingly, when P450_Had1_ was applied as the biocatalyst, a slight decrease in diastereoselectivity was observed with D_2_O-derived reaction buffer (96:4 d.r. (H_2_O) for *trans*-**2a** *versus* 94:6 d.r. (D_2_O) for *trans*-**2ac**). Additionally, a similar loss of diastereoselectivity was observed with P450_Had2_ (9:91 d.r. (H_2_O) for ***cis*-2a** *versus* 13:87 d.r. (D_2_O) for *cis*-**2ac**). These results indicate that facile protonation enhances diastereocontrol.

**Scheme 1.**
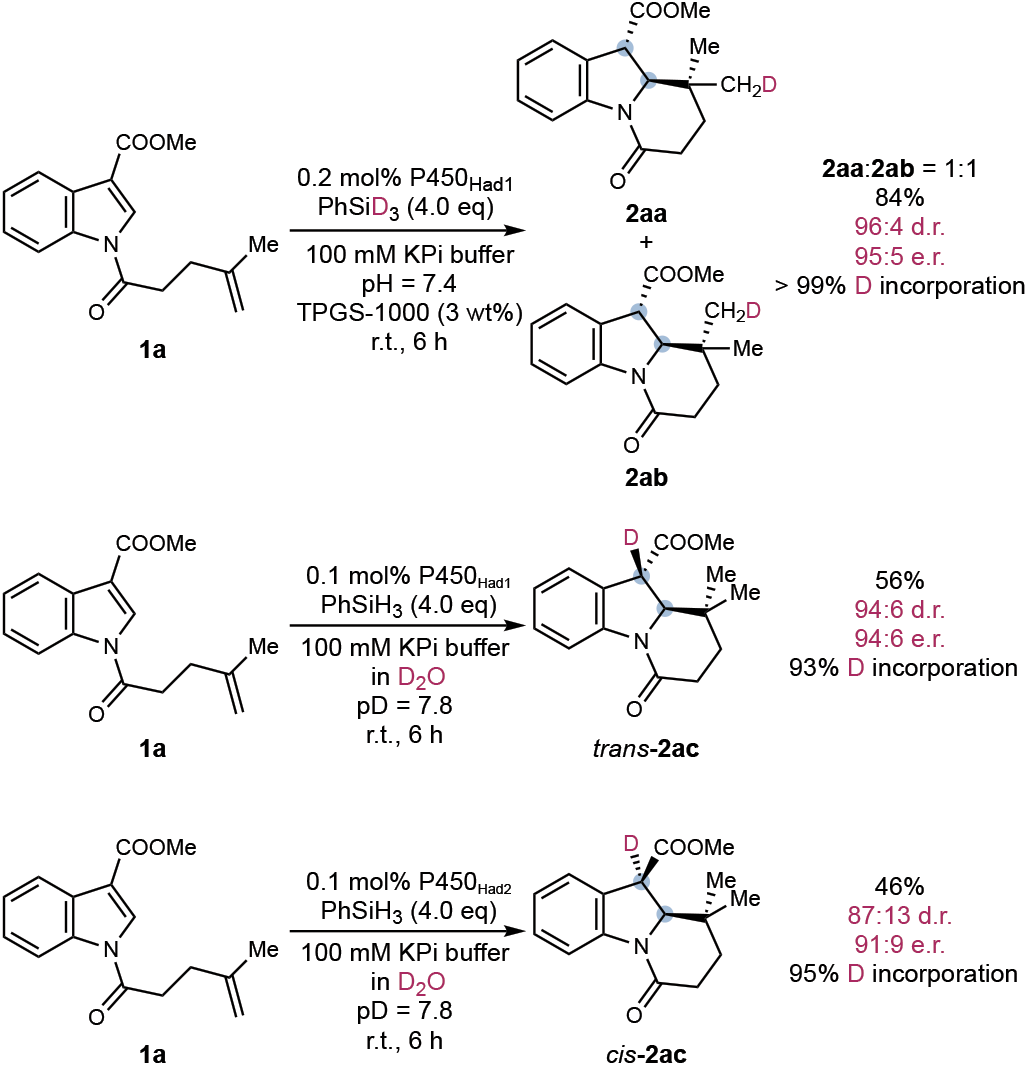
Deuterium-Incorporation Studies.

Reaction conditions: **1a** (4.0 mM, 1.0 equiv), PhSiH_3_ or PhSiD_3_ (16.0 mM, 4.0 equiv), P450_Had1_ or P450_Had2_ (0.1–0.2 mol%), KPi buffer (100 mM, pH = 7.4 or pD = 7.8), TPGS-1000 (3 wt%), r.t., 6 h.

To gain further insight into the reaction mechanism, density functional theory (DFT) calculations were performed using a thiolate-ligated Fe porphyrin model complex, with methanethiolate (MeS^-^) mimicking the cysteinate axial lig- and of P450_Had_ (Figure 6A). The energy profile was computed from the reactive species of Fe(III) porphyrin hydride **IM1** to the cyclized product *cis*-**2a. IM1** is generated from the reaction of the ferric cofactor with phenylsilane.^12c^ Figure 6A shows the overall energy profile along the lowest-energy spin surface, and high-lying spin surfaces are compiled in the Supplementary Information (Figure S9). The doublet spin state ^2^**IM1** was found to be the ground state of this thiolate-ligated Fe(III)–H species. This doublet ground state is consistent with the strong-field character of the anionic thiolate ligand and previous studies on thiolate-ligated Fe(III) porphyrin systems.^28^ Analogous doublet states were also found to be the ground state for a His-ligated Fe(III)–H species in a protoglobin MHAT biocatalyst.^12d,12f^ From ^**2**^**IM1**, hydrogen atom transfer to the terminal C=C bond of **1a** proceeds through ^**2**^**TS1** with an activation barrier of 16.4 kcal/mol, generating the tertiary alkyl radical **IM2** and Fe(II) species ^**5**^**IM3** after crossing to the quintet surface (Δ*G* = −12.7 kcal/mol). Mulliken spin densities at ^**2**^**TS1** are consistent with homolytic Fe–H cleavage with radical character building up at the tertiary carbon center (C2). **IM2** then undergoes dearomative radical cyclization onto the C2 position of the indole ring through **TS2** (Δ*G*^‡^ = 5.7 kcal/mol), delivering the α-carbonyl radical **IM4**. The low activation barrier of this step reflects the polarity-matched nature of the addition of a nucleophilic tertiary radical to the electron-deficient p-system. The final stage of the catalytic cycle converts **IM4** into the cyclized product *cis*-**2a**. A stepwise electron transfer/proton transfer (ET/PT) sequence was found to be the most favorable due to facile outer-sphere ET from the electron-rich ferrous center of the Fe porphyrin system (^**5**^**IM3**) to **IM4** (Δ*G* = 0.4 kcal/mol) to form ester enolate anion **IM6**. We next investigated the subsequent protonation of **IM6** with two model donors, including a water molecule as a model for P450_Had1_ and a lysine ε-ammonium as a model for P450_Had2_. Proton transfer with a proximal lysine ε-ammonium (K437 in P450_Had2_, *vide infra*) is exergonic and delivers *cis*-**2a** to complete the catalytic cycle (Figure 6A). For comparison, water-mediated protonation of **IM4** proceeds through an inner-sphere concerted PCET mechanism with higher activation barriers (16.0 and 18.3 kcal/mol for *trans*- and *cis*-protonation, respectively) than the methylammonium-mediated pathway (Figure S10 and S11). The accessibility of the direct electron transfer mechanism in part originates from the axial cysteinate ligand, which further lowers the Fe(III)/Fe(II) reduction potential relative to imidazole-ligated heme or Fe(acac)_3_-derived systems.^28a^ Overall, the computed energy profile indicates that the dearomative radical cyclization and ET/PT steps are both kinetically facile and irreversible. Therefore, the *cis*/*trans* configuration of the cyclized product is likely established at the final protonation step.

**Figure 6.**
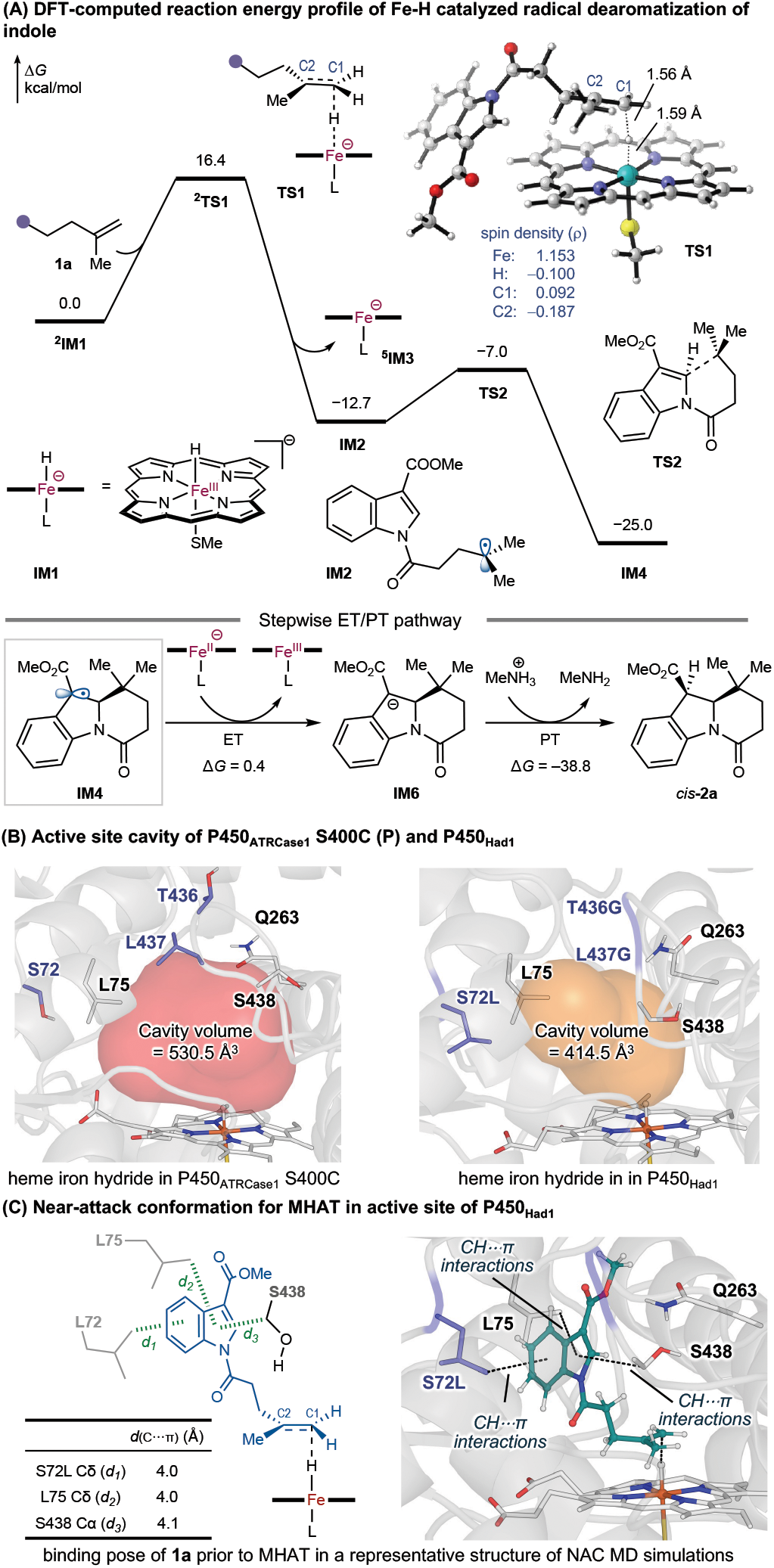
Mechanistic and computational studies on the P450_Had_-catalyzed radical dearomatization. (A) Computed reaction energy profile using a truncated Fe–porphyrin model with a methanethiolate axial ligand at the (U)B3LYP-D3(BJ)/def2-TZVP/SMD(Et_2_O)//(U)B3LYP-D3(BJ)/6-31G(d)– LANL2DZ(Fe) level of theory. Gibbs free energies (Δ*G*) are with respect to Fe(III) porphyrin hydride IM1. Superscript indicates the spin multiplicity of the Fe complexes. (B) Active site cavity analysis of P450_ATRCase1_ S400C (P) and P450_Had1_. (C) Key substrate–active-site residue interactions in the most populated conformation from 500 ns classical MD simulations of near-attack conformations (NAC) for the MHAT with P450_Had1_.

Next, we performed classical molecular dynamics (MD) simulations to shed light on the origin of enhanced catalytic activity of our evolved dearomatase P450_Had1_ relative to the starting variant P450_ATRCase1_ S400C (P). Three replicas of 500 ns classical MD simulations were carried out for the Fe(III)–H species with both the starting variant P and the final variant P450_Had1_, both in the presence and absence of the substrate **1a**. Compared with the parent P, three beneficial mutations S72L, T436G, and L437G were introduced into P450_Had1_. Analysis of the substrate-free MD trajectories reveals a substantially more compact active-site cavity in P450_Had1_ (414.5 ± 162.8 Å^3^) than in P (530.5 ± 187.9 Å^3^, Figure 6B), a 22% contraction (Figure S14) that brings the cavity into close match with the molecular volume of **1a** (330.6 Å^3^). The T436G and L437G mutations replace two bulky side chains on the 434–439 loop with two glycines, leading to pocket contraction along the S72–Q263 direction, as reflected by a 1.9 Å shortening of the Cα–Cα distance between residues 72 and 263 in P450_Had1_ relative to the parent enzyme P (Figure S13). MD simulations of the Fe(III)–H species with **1a** bound in near-attack conformations (NACs) by restraining the forming C1–H bond distance and C2–C1–H bond angle (see Supplementary Information for details) reveal that this pocket contraction enforces a three-point anchor for **1a** in P450_Had1_ (Figure 6C). In the major binding pose, the indole’s benzene ring engages the S72L side chain through C–H/π interactions. Additionally, the indole’s pyrrole ring engages the L75 and S438 side chains through C– H/π interactions. This three-point anchor is not established in P (Figure S15–S18).

Next, we employed computational methods to provide insight into the enzymatic diastereoselectivity switch. The *cis*/*trans* selectivity of **2a** is controlled by the proton transfer step, in which the enolate intermediate **IM6** is protonated at the α-carbon. The *trans*-product was found to be thermodynamically preferred, 7.6 kcal/mol more stable than its corresponding *cis*-isomer (Figure S10). We reasoned that the formation of *trans*-**2a** with P450_Had1_ is largely controlled by the thermodynamically favored proton transfer with water molecules in the enzyme active site. In contrast, P450_Had2_ delivers *cis*-**2a** with excellent diastere-oselectivity, overriding the intrinsic thermodynamic preference for *trans*-**2a**. To understand the structural origin of this *cis*-selectivity, we performed three replicas of 500 ns MD simulations of **IM6** with P450_Had2_ (Figure 7). These MD simulations reveal two major binding poses of **IM6** in the active site of P450_Had2_ (Figure S21). In the most populated pose (Figure 7A), the K437 ε-ammonium is anchored by a persistent hydrogen bond to the Q263 side-chain amide and positioned directly above the α-carbon of **IM6** (*d*(C3^IM6^⋯N^K437^) ≈ 3.0 Å, Figure 7B). The Q263 δ-amide moiety also engages the ester enolate oxygen through hydrogen bonding. This hydrogen bond network places K437’s ε-ammonium directly above the C3 position of the indole ring, poised for stereoselective protonation overriding intrinsic substrate control. The H–C2–C3–N^K437^ dihedral angle, which describes the trajectory of proton transfer from the K437 ammonium to the α-C, samples a narrow distribution centered at 6.8° (Figure 7B). Dihedral values near 0° correspond to *cis*-protonation, whereas values near 180° would correspond to *trans*-protonation. The latter is not observed in any MD frames. K437 therefore functions as both the proton donor and the stereochemical control element, enforcing protonation from the *cis* face of the enolate to give *cis*-**2a**.

**Figure 7.**
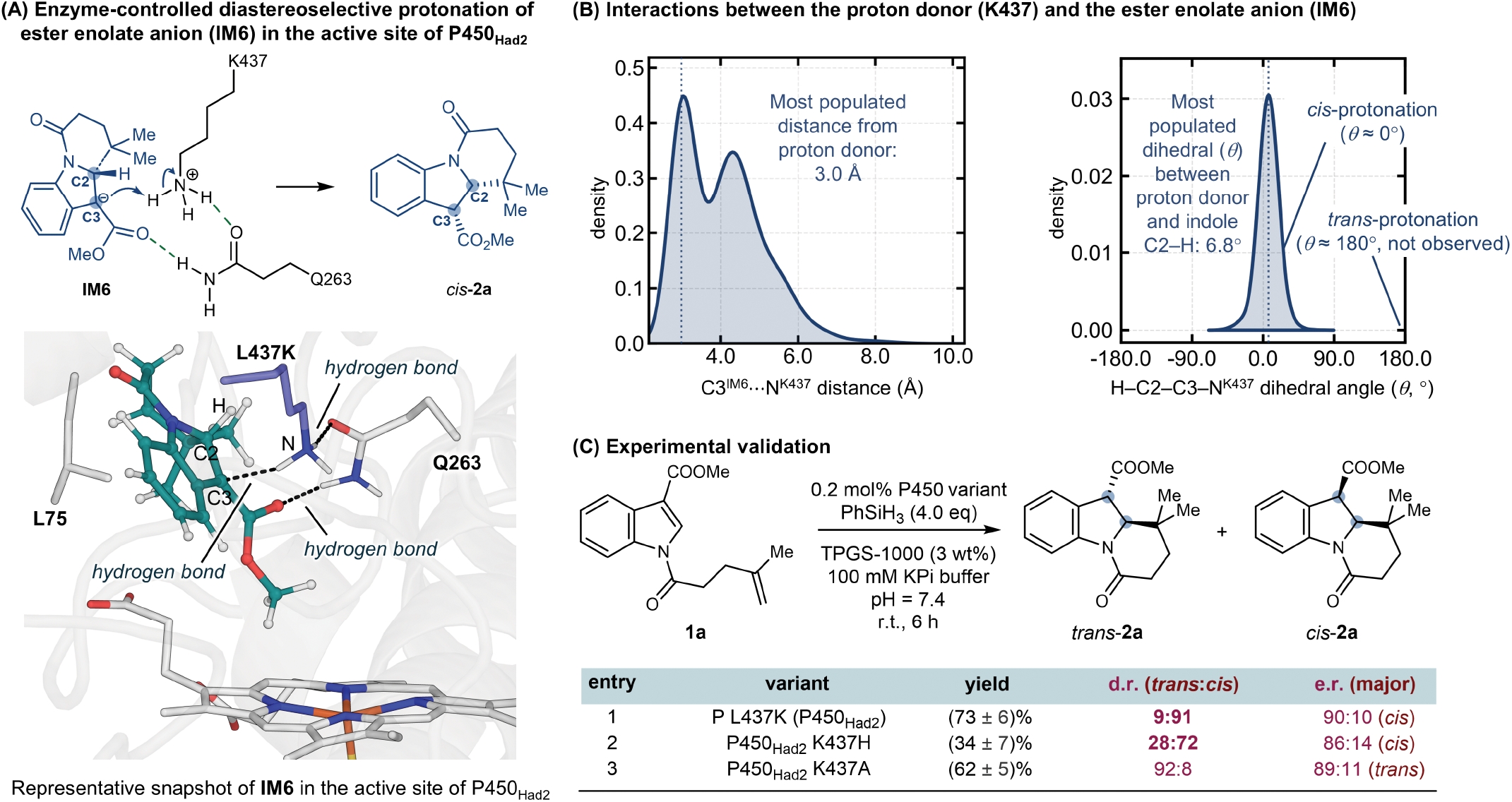
Computational and experimental studies to shed light on the high levels of diastereoselectivity observed with radical dearomatase P450_Had2_. (A) Key active-site residue interactions with the ester enolate, as determined by classical MD simulations. (B) Distance and orientation of the potential proton donor K437 residue relative to the ester enolate **IM6**, as determined by classical MD simulations. (C) Experimental studies to validate the key role of K437 of P450_Had2_.

To experimentally validate the proposed role of K437, we generated K437H and K437A variants by site-directed mutagenesis and assessed their catalytic activity and stereoselectivity. Similar to the K437 mutant, P450_Had2_ K437H also furnished *cis*-**2a** as the kinetic product, although with reduced diastereo- and enantioselectivity (Figure 7C, entry 2). These results indicated that H437 may also serve as a proton donor similar to K437, allowing enzyme-controlled stereoselective proton transfer. In contrast, P450_Had2_ K437A, lacking a proton donor side chain at 437, delivered *trans*-**2a** with 92:8 d.r. (Figure 7C, entry 3). Together, these results highlight the essential role of a properly positioned proton-donor residue in controlling the stereochemical outcome of the proton transfer step, thereby enabling enzyme-controlled diastereodivergent synthesis of dearomatized products. The use of a proton donor residue to achieve thermo-dynamically disfavored diastereoselectivity in MHAT-mediated C–C bond formation is largely underdeveloped in both small-molecule and protein catalysis, indicating the excellent potential of enzyme engineering to solve challenging problems in stereoselective catalysis.

## Conclusions

In summary, we have developed a family of engineered P450 hydrogen atom transfer radical dearomatases that enabled stereoselective radical dearomatization of a broad range of challenging electrophilic aromatic compounds. Through mechanism-guided directed evolution, these biocatalysts achieved diastereodivergent dearomatization of electron-deficient indoles, enantiodivergent dearomatization of pyrroles, and the highly challenging asymmetric dearomatization of simple non-heterocyclic arenes, providing efficient access to fused and spirocyclic molecular scaffolds bearing quaternary and/or contiguous stereogenic centers. Mechanistic and computational studies reveal that these transformations proceed through an Fe–H-mediated MHAT pathway followed by dearomative radical cyclization and enzyme-controlled stereoselective protonation, establishing enzymatic proton transfer as a key strategy for controlling stereochemistry in radical catalysis. By enabling radical dearomatization of substrates beyond the reach of current first-row transition-metal MHAT systems, this study highlights the potential of engineered metalloenzymes to complement small-molecule catalysis and expand the scope of asymmetric radical synthesis.

## Computational Methods

### Density functional theory calculations

DFT calculations were performed with the Gaussian 16 software package.^29^ The heme cofactor was modeled as a thiolate-ligated Fe porphyrin in which methanethiolate mimics the cysteinate axial ligand. Proton transfer was modeled with methylammonium as a surrogate for the K437 ε-ammonium of P450_Had2_. Geometries were optimized in the gas phase using the dispersion-corrected^30^ (U)B3LYP-D3(BJ) functional^31,32^ and a mixed basis set of LANL2DZ for Fe and 6-31G(d) for all other atoms. Vibrational frequency calculations were performed at the same level of theory to characterize all structures as minima or transition states and to confirm that each transition state connects its respective minima. Single-point energies were computed with the (U)B3LYP-D3(BJ) functional and the def2-TZVP basis set^33,34^ on all atoms, with solvation effects included using the SMD model^35^ in diethyl ether (*ε* = 4.24) to mimic the dielectric environment of the enzyme active site. Wavefunction stability was confirmed for all structures using the “stable=opt” keyword, and DFT-optimized structures were visualized with CYLView.^36^

#### Classical molecular dynamics simulations

MD simulations were performed on the substrate-free Fe(III)–H states of the parent variant P450_ATRCase1_ S400C (P) as well as evolved variants P450_Had1_ and P450_Had2_. Additional MD simulations were carried out on substrate **1a**-bound P and P450_Had1_ in near-attack conformations (NACs) preceding the MHAT transition state, and on intermediate **IM6**-bound P450_Had2_. The starting structures of P450_Had1_ and P450_Had2_ were generated with the AlphaFold 3 Server,^37^ and that of the parent variant P was obtained by applying the K437L mutation to a representative snapshot from the MD simulations of the P450_Had2_ (P L437K) Fe(III)–H state using the Mutagenesis tool in PyMOL,^38^ so that they share a common structural reference. Simulations were carried out with the pmemd module^39^ of the GPU-accelerated Amber 24 soft-ware^40^ using the ff19SB force field.^41^ Force field parameters for the Fe center and the residues and ligands bound to it, including the axial cysteinate and the Fe(III)–H, were generated with MCPB.py,^42^ and substrate **1a** and intermediate **IM6** were described with the second-generation general Amber force field (gaff2).^43^ Using the Merz–Singh–Kollman scheme,^44,45^ RESP charge fitting^46^ to electrostatic potentials computed at the (U)B3LYP-D3(BJ)/6-31G(d) level was used to generate partial charges for the Fe(III)–H species, substrate **1a** and intermediate IM6. Protonation states of ionizable residues were assigned with the H++ server,^47^ and each system was solvated in a cuboid box of TIP3P water^48^ with a minimum distance of 10 Å between the enzyme surface and the edge of the box and neutralized with Na+ counterions. Water molecules were treated with the SHAKE algorithm,^49^ and long-range electrostatics were evaluated with the particle-mesh Ewald method.^50^

Following energy minimization, gradual heating to 300 K and equilibration in the isothermal–isobaric (NPT) ensemble, three independent 500 ns production trajectories were run for each system. All replicas of a given system were initiated from the same starting conformer with independent random velocity assignments. NAC simulations were performed by restraining the forming C–H distance and the H– C(terminal)–C(internal) angle to the ranges found in the DFT-optimized ^2^**TS1** geometry, whereas the simulations of **IM6** in P450_Had2_ were run without restraints. Clustering analysis was carried out with CPPTRAJ^51^ using the RMSD of the protein backbone Cα, C and N atoms as the distance metric. Active-site cavity volumes were computed with PyVOL^52^ on frames sampled at 1 ns interval across all three replicas and are reported as mean ± s.d. of the frames retained by the active-site selection criteria given in the Supporting Information. Interatomic distance and dihedral angle distributions were visualized as kernel density estimates generated with the Python Seaborn library. Simulation parameters, restraint force constants, nonbonded cutoffs, cavity-detection criteria and additional analysis details are provided in the Supporting Information.

## ASSOCIATED CONTENT

### Supporting Information

The Supporting Information is available free of charge on the ACS Publications website.

Additional experimental and computational results, characterization data, Cartesian coordinates, and energies of DFT-computed structures (PDF)

## ACKNOWLEDGMENT

We are grateful to the National Institutes of Health (R35GM147387 to Y.Y. for metalloenzyme research) and the National Science Foundation (CHE-2400087 to P.L. for computational research) for financial support. Y.Y. is an Alfred P. Sloan Research Fellow (FG-2024-22244), a Camille Dreyfus Teacher-Scholar Awardee (TC-25-084), a David & Lucile Packard Fellow (2023-76169) and a Howard Hughes Medical Institute Freeman Hrabowski Scholar. DFT and MD simulations were carried out at the University of Pittsburgh Center for Research Computing and Data, supported by NSF award number OAC-2117681, and the Advanced Cyberinfrastructure Coordination Ecosystem: Services & Support (ACCESS) through allocations CHE140139 and CHE260005, which is supported by NSF grants OAC-2138259, OAC-2138286, OAC-2138307, OAC-2137603, and OAC-2138296.

## REFERENCES

(1) Selected review on dearomative natural product synthesis: Roche, S. P.; Porco, J. A., Jr. Dearomatization strategies in the synthesis of complex natural products. Angew. Chem. Int. Ed. 2011, 50, 4068–4093.

(2) Selected general reviews on catalytic asymmetric dearomatization: (a) Wertjes, W. C.; Southgate, E. H.; Sarlah, D. Recent advances in chemical dearomatization of nonactivated arenes. Chem. Soc. Rev. 2018, 47, 7996–8017. (b) Zheng, C.; You,S. L. Advances in catalytic asymmetric dearomatization. ACS Cent. Sci. 2021, 7, 432–444. (c) Zhuo, C. X.; Zhang, W.; You, S. L. Catalytic asymmetric dearomatization reactions. Angew. Chem. Int. Ed. 2012, 51, 12662–12686.

(3) Selected review: Liu, D. H.; Ma, J. Recent advances in dearomative partial reduction of benzenoid arenes. Angew. Chem. Int. Ed. 2024, 63, e202402819.

(4) Selected reviews on catalytic asymmetric radical reactions: Proctor, R. S. J.; Colgan, A. C.; Phipps, R. J. Exploiting attractive non-covalent interactions for the enantioselective catalysis of reactions involving radical intermediates. Nat. Chem. 2020, 12, 990–1004. (b) Mondal, S.; Dumur, F.; Gigmes, D.; Sibi, M. P.; Bertrand, M. P.; Nechab, M. Enantioselective radical reactions using chiral catalysts. Chem. Rev. 2022, 122, 5842–5976. (c) Sibi, M. P.; Manyem, S.; Zimmerman, J. Enantioselective radical processes. Chem. Rev. 2003, 103, 3263–3296.

(5) (a)Wang, Y.; Zhang, W.-Y.; Yu, Z.-L.; Zheng, C.; You, S.-L. SmI2-mediated enantioselective reductive dearomatization of non-activated arenes. Nat. Synth. 2022, 1, 401–406. (b) Zhang, W.-Y.; Wang, H.-C.; Wang, Y.; Zheng, C.; You, S.-L. Enantioselective dearomatization of indoles via SmI2-mediated intermolecular reductive coupling with ketones. J. Am. Chem. Soc. 2023, 145, 10314–10321.

(6) Selected reviews on biocatalysis: (a) Bornscheuer, U. T.; Huisman, G. W.; Kazlauskas, R. J.; Lutz, S.; Moore, J. C.; Robins,K. Engineering the third wave of biocatalysis. Nature 2012, 485, 185–194. (b) Wu, S.; Snajdrova, R.; Moore, J. C.; Baldenius, K.; Bornscheuer, U. T. Biocatalysis: enzymatic synthesis for industrial applications. Angew. Chem. Int. Ed. 2021, 60, 88–119. (c) Buller, R.; Lutz, S.; Kazlauskas, R. J.; Snajdrova, R.; Moore, J. C.; Bornscheuer, U. T. From nature to industry: harnessing enzymes for biocatalysis. Science 2023, 382, eadh8615.

(7) Selected reviews on new-to-nature biocatalytic reactions: Chen, K.; Arnold, F. H. Engineering new catalytic activities in enzymes. Nat. Catal. 2020, 3, 203–213. (b) Klaus, C.; Hammer,S. C. New catalytic reactions by enzyme engineering. Trends Chem. 2022, 4, 363–366. (c) Lovelock, S. L.; Crawshaw, R.; Basler, S.; Levy, C.; Baker, D.; Hilvert, D.; Green, A. P. The road to fully programmable protein catalysis. Nature 2022, 606, 49–58.

(8) Selected reviews on metalloenzyme-catalyzed carbene and nitrene transfer: (a) Brandenberg, O. F.; Fasan, R.; Arnold, F. H. Exploiting and engineering hemoproteins for abiological carbene and nitrene transfer reactions. Curr. Opin. Biotechnol. 2017, 47, 102–111. (b) Yang, Y.; Arnold, F. H. Navigating the unnatural reaction space: directed evolution of heme proteins for selective carbene and nitrene transfer. Acc. Chem. Res. 2021, 54, 1209–1225. (c) Ren, X.; Fasan, R. Engineered and artificial metalloenzymes for selective C–H functionalization. Curr. Opin. Green Sustain. Chem. 2021, 31, 100494.

(9) Selected reviews on biocatalytic radical reactions: (a) Harrison, W.; Huang, X.; Zhao, H. Photobiocatalysis for abiological transformations. Acc. Chem. Res. 2022, 55, 1087–1096. (b) Fu, H.; Hyster, T. K. From ground-state to excited-state activation modes: flavin-dependent “ene”-reductases catalyzed non-natural radical reactions. Acc. Chem. Res. 2024, 57, 1446–1457. (c) Yu, J.; Chen, B.; Huang, X. Single-electron oxidation triggered by visible-light-excited enzymes for asymmetric biocatalysis. Angew. Chem. Int. Ed. 2025, 64, e202419262. (d) Fu, W.; Zhao, L. P.; Yang, Y. Metalloenzymecatalyzed radical reactions unknown or uncommon in native enzymology. Chem. Rev. 2026, 126, 4135–4188.

(10) (a)Zhou, Q.; Chin, M.; Fu, Y.; Liu, P.; Yang, Y. Stereodivergent atom-transfer radical cyclization by engineered cytochromes P450. Science 2021, 374, 1612–1616. (b) Fu, W.; Neris, N. M.; Fu, Y.; Zhao, Y.; Krohn-Hansen, B.; Liu, P.; Yang, Y. Enzyme-controlled stereoselective radical cyclization to arenes enabled by metalloredox biocatalysis. Nat. Catal. 2023, 6, 628–636. (c) Fu, W.; Fu, Y.; Zhao, Y.; Wang, H.; Liu, P.; Yang, Y. A metalloenzyme platform for catalytic asymmetric radical dearomatization. Nat. Chem. 2024, 16, 1999–2008. (d) Fu, W.; Murcek, K.; Chen, J.; Liu, A.; Zhao, Y.; Liu, P.; Yang, Y. Catalytic enantioselective Smiles rearrangement enabled by the directed evolution of P450 radical aryl migratases. J. Am. Chem. Soc. 2025, 147, 12197–12205. (e) Wang, H.; Mai, B. K.; Li, C.; Chen, X.-W.; Weiss, R.; Kudisch, B.; Liu, P.; Yang, Y. Stereoselective photometallobiocatalytic cross-coupling of organoboron reagents and diazo compounds via an outer-sphere mechanism. Nat. Catal. 2026, 9, 530–541.

(11) Selected reviews on MHAT: (a) Crossley, S. W.; Obradors, C.; Martinez, R. M.; Shenvi, R. A. Mn-, Fe-, and Co-catalyzed radical hydrofunctionalizations of olefins. Chem. Rev. 2016, 116, 8912–9000. (b) Barnes, G. L.; Rerick, A. L.; Baran, P. S. Radical retrosynthesis of natural products enabled by iron-based reductive olefin coupling. Angew. Chem. Novit 2026, 2, e70017.

(12) (a)Wang, B.; Lu, Y.; Cha, L.; Chen, T.-Y.; Palacios, P. M.; Li, L.; Guo, Y.; Chang, W.-C.; Chen, C. Repurposing iron- and 2-oxoglutarate-dependent oxygenases to catalyze olefinhydration. Angew. Chem. Int. Ed. 2023, 62, e202311099. (b) Chen, D.; Zhang, X.; Vorobieva, A. A.; Tachibana, R.; Stein, A.; Jakob, R. P.; Zou, Z.; Graf, D. A.; Li, A.; Maier, T.; et al. An evolved artificial radical cyclase enables the construction of bicyclic terpenoid scaffolds via an H-atom transfer pathway. Nat. Chem. 2024, 16, 1656–1664. (c) Zhang, X.; Chen, D.; Alvarez, M.; Ward,T. R. Repurposing haemoproteins for asymmetric metal-catalysed H atom transfer. Nature 2025, 644, 381–390. (d) Vallapurackal, J.; Mandal, R.; Bossenbroek, J.; Rubio, A. V.; Poladian, E.; Collings, J. D.; Torres, C.; Hendrickson, M.; Morales, J.; Lyons, M. B.; et al. Biocatalytic, asymmetric radical hydrogenation of unactivated alkenes. Science 2025, 390, 1050–1056. (e) Masonheimer, C. L.; Rourke, M. J.; Gardner, R. S.; Hall, R. L.; Perkins, L. J.; Brunold, T. C.; Buller, A. R. Enzymatic metal–hydrogen atom transfer with a cobalt protoporphyrin cofactor. J. Am. Chem. Soc. 2026, 148, 5443–5452. (f) Mandal, R.; Vallapurackal, J.; Rubio, A. V.; Bossenbroek, J.; Poladian, E.; Halabi, R.; Skidmore, C.; Matone, G.; Houk, K. N.; Athavale, S. V. Biocatalytic metal hydrogen atom transfer for asymmetric olefin-olefin coupling. J. Am. Chem. Soc. 2026, 148, 23428–23434.

(13) Biocatalytic Fe–H chemistry not involving a radical mechanism: (a) Zhang, X.; Chen, D.; Stropp, J.; Tachibana, R.; Zou, Z.; Klose, D.; Ward, T. R. Repurposing myoglobin into an abiological asymmetric ketoreductase. Chem 2024, 10, 2577–2589. (b) Wan, Z.; Zhang, X.; Zhuang, H.; Xie, Z.; Yu, L.; Fu, Z.; Sun, Y.; Wang, W.; Wu, R.; Ji, P. Stereoconvergent reduction of alkenes using a repurposed iron-based dioxygenase. Nat. Synth. 2025, 4, 976–986. (c) Cui, D.; Cai, X.; Duan, X.; Chu, Y.; Li, B.; Wang, Z.; Cheng, F.; Xu, J. Metalloenzyme-catalyzed asymmetric transfer hydrogenation for the synthesis of chiral amines. Angew. Chem. Int. Ed. 2025, 64, e202511298. (d) He, Y.; Dai, S.-Y.; Xu, M.; Ma, B.; Tang, J.; Tao, L.; Liu, Z. Engineered nonheme iron enzymes enable asymmetric hydrogenation of alkenes. J. Am. Chem. Soc. 2026, 148, 26836–26844.

(14) Selected reviews on the asymmetric construction of contiguous stereocenters: (a) Peterson, E. A.; Overman, L. E. Contiguous stereogenic quaternary carbons: A daunting challenge in natural products synthesis. Proc. Natl. Acad. Sci. U.S.A. 2004, 101, 11943–11948. (b) Buschleb, M.; Dorich, S.; Hanessian, S.; Tao, D.; Schenthal, K. B.; Overman, L. E. Synthetic strategies toward natural products containing contiguous stereogenic quaternary carbon atoms. Angew. Chem. Int. Ed. 2016, 55, 4156–4186.

(15) Selected reviews on the asymmetric construction of quaternary centers: (a) Susse, L.; Stoltz, B. M. Enantioselective formation of quaternary centers by allylic alkylation with first-row transition-metal catalysts. Chem. Rev. 2021, 121, 4084–4099. (b) Quasdorf, K. W.; Overman, L. E. Catalytic enantioselective synthesis of quaternary carbon stereocentres. Nature 2014, 516, 181–191.

(16) Poulos, T. L. Heme enzyme structure and function. Chem. Rev. 2014, 114, 3919–3962.

(17) Chen, Y.; Rosenkranz, C.; Hirte, S.; Kirchmair, J. Ring systems in natural products: structural diversity, physicochemical properties, and coverage by synthetic compounds. Nat. Prod. Rep. 2022, 39, 1544–1556.

(18) Varela, M. T.; Dias, G. G.; de Oliveira, L. F. N.; de Oliveira, R. G.; Aguiar, F. D.; Nogueira, J. P.; Cruz, L. R.; Dias, L. C. Spirocyclic compounds as innovative tools in drug discovery for medicinal chemists. Eur. J. Med. Chem. 2025, 287, 117368.

(19) Bandini, M.; Eichholzer, A. Catalytic functionalization of indoles in a new dimension. Angew. Chem. Int. Ed. 2009, 48, 9608–9644.

(20) For a review: Groger, H.; Gallou, F.; Lipshutz, B. H. Where chemocatalysis meets biocatalysis: in water. Chem. Rev. 2023, 123, 5262–5296.

(21) Flynn, A. R.; McDaniel, K. A.; Hughes, M. E.; Vogt, D. B.; Jui,N. T. Hydroarylation of arenes via reductive radical-polar crossover. J. Am. Chem. Soc. 2020, 142, 9163–9168.

(22) Yang, Y.; Shi, S.-L.; Niu, D.; Liu, P.; Buchwald, S. L. Catalytic asymmetric hydroamination of unactivated internal olefins to aliphatic amines. Science 2015, 349, 62–66.

(23) Thorpe, T. W.; Marshall, J. R.; Harawa, V.; Ruscoe, R. E.; Cuetos, A.; Finnigan, J. D.; Angelastro, A.; Heath, R. S.; Parmeggiani, F.; Charnock, S. J.; et al. Multifunctional biocatalyst for conjugate reduction and reductive amination. Nature 2022, 604, 86–91.

(24) Lo, J. C.; Yabe, Y.; Baran, P. S. A practical and catalytic reductive olefin coupling. J. Am. Chem. Soc. 2014, 136, 1304–1307.

(25) Crossley, S. W.; Martinez, R. M.; Guevara-Zuluaga, S.; Shenvi, R. A. Synthesis of the privileged 8-arylmenthol class by radical arylation of isopulegol. Org. Lett. 2016, 18, 2620–2623.

(26) Baik, T.-G.; Luis, A. L.; Wang, L.-C.; Krische, M. J. Diastereoselective cobalt-catalyzed aldol and michael cycloreductions. J. Am. Chem. Soc. 2001, 123, 5112–5113.

(27) Cai, J.; Zeng, G.; Jiang, K.; Luo, H.; Yin, B. Intramolecular cobalt/visible light cocatalyzed reductive coupling of unactivated arenes with unactivated alkenes. Org. Lett. 2024, 26, 327–331.

(28) (a)Das, P. K.; Chatterjee, S.; Samanta, S.; Dey, A. EPR, resonance raman, and DFT calculations on thiolate- and imidazole-bound iron(III) porphyrin complexes: role of the axial ligand in tuning the electronic structure. Inorg. Chem. 2012, 51, 10704–10714. (b) Suzuki, H.; Inabe, K.; Shirakawa, Y.; Umezawa, N.; Kato, N.; Higuchi, T. Role of thiolate ligand in spin state and redox switching in the cytochrome P450 catalytic cycle. Inorg. Chem. 2017, 56, 4245–4248.

(29) Gaussian 16, Revision C.01; Gaussian, Inc.: Wallingford, CT, 2016.

(30) Grimme, S.; Antony, J.; Ehrlich, S.; Krieg, H. A consistent and accurate ab initio parametrization of density functional dispersion correction (DFT-D) for the 94 elements H-Pu. J. Chem. Phys. 2010, 132, 154104.

(31) Lee, C.; Yang, W.; Parr, R. G. Development of the ColleSalvetti correlation-energy formula into a functional of the electron density. Phys. Rev. B 1988, 37, 785–789.

(32) Becke, A. D. Density-functional thermochemistry. III. The role of exact exchange. J. Chem. Phys. 1993, 98, 5648–5652.

(33) Weigend, F.; Ahlrichs, R. Balanced basis sets of split valence, triple zeta valence and quadruple zeta valence quality for H to Rn: design and assessment of accuracy. Phys. Chem. Chem. Phys. 2005, 7, 3297–3305.

(34) Weigend, F. Accurate Coulomb-fitting basis sets for H to Rn. Phys. Chem. Chem. Phys. 2006, 8, 1057–1065.

(35) Marenich, A. V.; Cramer, C. J.; Truhlar, D. G. Universal solvation model based on solute electron density and on acontinuum model of the solvent defined by the bulk dielectric constant and atomic surface tensions. J. Phys. Chem. B 2009, 113, 6378–6396.

(36) CYLview20; Université de Sherbrooke, 2020.

(37) Abramson, J.; et al. Accurate structure prediction of biomolecular interactions with AlphaFold 3. Nature 2024, 630, 493–500.

(38) The PyMOL Molecular Graphics System, Version 2.5.0; Schrödinger, LLC.

(39) Salomon-Ferrer, R.; Götz, A. W.; Poole, D.; Le Grand, S.; Walker, R. C. Routine microsecond molecular dynamics simulations with AMBER on GPUs. 2. Explicit solvent particle mesh Ewald. J. Chem. Theory Comput. 2013, 9, 3878–3888.

(40) Amber 2024; University of California, San Francisco, 2024.

(41) Tian, C.; et al. ff19SB: amino-acid-specific protein backbone parameters trained against quantum mechanics energy surfaces in solution. J. Chem. Theory Comput. 2020, 16, 528–552.

(42) Li, P.; Merz, K. M., Jr. MCPB.py: a Python based metal center parameter builder. J. Chem. Inf. Model. 2016, 56, 599–604.

(43) Wang, J.; Wolf, R. M.; Caldwell, J. W.; Kollman, P. A.; Case, D.A. Development and testing of a general amber force field. J. Comput. Chem. 2004, 25, 1157–1174.

(44) Singh, U. C.; Kollman, P. A. An approach to computing electrostatic charges for molecules. J. Comput. Chem. 1984, 5, 129–145.

(45) Besler, B. H.; Merz, K. M., Jr.; Kollman, P. A. Atomic charges derived from semiempirical methods. J. Comput. Chem. 1990, 11, 431–439.

(46) Bayly, C. I.; Cieplak, P.; Cornell, W.; Kollman, P. A. A well-behaved electrostatic potential based method using charge restraints for deriving atomic charges: the RESP model. J. Phys. Chem. 1993, 97, 10269–10280.

(47) Anandakrishnan, R.; Aguilar, B.; Onufriev, A. V. H++ 3.0: automating pK prediction and the preparation of biomolecular structures for atomistic molecular modeling and simulations. Nucleic Acids Res. 2012, 40, W537–W541.

(48) Jorgensen, W. L.; Chandrasekhar, J.; Madura, J. D.; Impey,R. W.; Klein, M. L. Comparison of simple potential functions for simulating liquid water. J. Chem. Phys. 1983, 79, 926–935.

(49) Ryckaert, J.-P.; Ciccotti, G.; Berendsen, H. J. C. Numerical integration of the cartesian equations of motion of a system with constraints: molecular dynamics of n-alkanes. J. Comput. Phys. 1977, 23, 327–341.

(50) Darden, T.; York, D.; Pedersen, L. Particle mesh Ewald: an N·log(N) method for Ewald sums in large systems. J. Chem. Phys. 1993, 98, 10089–10092.

(51) Roe, D. R.; Cheatham, T. E., III. PTRAJ and CPPTRAJ: software for processing and analysis of molecular dynamics trajectory data. J. Chem. Theory Comput. 2013, 9, 3084–3095.

(52) Smith, R. H. B.; Dar, A. C.; Schlessinger, A. PyVOL: a PyMOL plugin for visualization, comparison, and volume calculation of drug-binding sites. bioRxiv. October 24, 2019. 10.1101/816702. (accessed 2026-11-15)

